# A novel neural stem cell-derived immunocompetent mouse model of glioblastoma for preclinical studies

**DOI:** 10.1101/2020.03.16.993196

**Authors:** Barbara Costa, Michael Fletcher, Pavle Boskovic, Ekaterina L. Ivanova, Tanja Eisemann, Sabrina Lohr, Lukas Bunse, Martin Löwer, Stefanie Burchard, Andrey Korshunov, Hai-Kun Liu, Michael Platten, Bernhard Radlwimmer, Peter Angel, Heike Peterziel

**Author notes:** **Contributions** B.C., P.A., and H.P. developed the study concept and design; B.C., M.F., P.B., E.I., T.E., S.L., L.B., M.L. conducted the experimental work and analyzed and interpreted the data; A.K. scored the immunohistochemistry of the tumors; B.C., P.B., M.F., P.A., and H.P. wrote the original draft; and all authors revised and approved the manuscript before submission. P.A. and H.P. equally contributed to senior authorship. **Corresponding authors** Peter Angel, Division of Signal Transduction and Growth Control, DKFZ-ZMBH Alliance, German Cancer Research Center (DKFZ), Im Neuenheimer Feld 280, 69120 Heidelberg, Germany; Barbara Costa, Division of Signal Transduction and Growth Control, DKFZ-ZMBH Alliance, German Cancer Research Center (DKFZ), Im Neuenheimer Feld 280, 69120 Heidelberg, Germany.

## Abstract

Glioblastomas are the most lethal tumors affecting the central nervous system in adults. Simple and inexpensive syngeneic *in vivo* models that closely mirror human glioblastoma, including interactions between tumor and immune cells, are urgently needed for deciphering glioma biology and developing more effective treatments. Here, we generated glioblastoma cell lines by repeated *in-vivo* passaging of cells isolated from a neural stem cell-specific Pten/p53 double-knockout genetic mouse brain tumor model. Transcriptome and genome analyses of the cell lines revealed molecular heterogeneity comparable to that observed in human glioblastoma. Upon orthotopic transplantation into syngeneic hosts they formed high-grade gliomas that faithfully recapitulated the histopathological features, invasiveness and myeloid cell infiltration characteristic of human glioblastoma. These features make our cell lines unique and useful tools to study multiple aspects of glioblastoma pathomechanism and test novel treatments, especially immunotherapies in syngeneic preclinical models.

## Introduction

Glioblastomas are the most prevalent malignant brain tumors in adults. These WHO grade IV tumors are defined by histopathologic features including mitotic figures, microvascular proliferation, necrosis and extensive infiltration of the brain parenchyma, which impairs complete surgical removal almost invariably leading to tumor relapse. Despite the presence of immune infiltration, typically constituted by myeloid cells, glioblastomas display a highly immunosuppressive tumor microenvironment with a low number of tumor-infiltrating lymphocytes^1^. At the molecular level 90% of glioblastomas harbor wildtype isocitrate dehydrogenase 1 and 2 genes (IDH1 and IDH2) distinguishing them from the remaining 10%, the so-called secondary glioblastomas, with a better prognosis^2^. The current treatment regimen of glioblastoma patients consists of maximal surgical resection combined with chemo- and radiotherapy; however, the median survival of about 15 months post diagnosis has not improved substantially in the recent decades, making glioblastoma one of the deadliest tumors and underscoring the importance of developing novel, more effective treatments^3^. Our knowledge of glioblastoma molecular pathomechanisms has greatly advanced in the past decade, and it has become widely accepted that IDH wildtype (IDH^wt^) glioblastomas can be sub-classified into three subtypes: classical, proneural, and mesenchymal^4^. Importantly, the corresponding subtype-specific RNA-expression-based signatures are also influenced by the composition of the tumor microenvironment, including stromal and immune cell populations^5^. Despite these advances, we are still lacking *in vivo* preclinical models that reflect the molecular and phenotypic heterogeneity of glioblastoma and recapitulate their complex interaction with the immune cells of the tumor microenvironment. In order to thoroughly understand the biology underlying glioblastoma development and for more predictive preclinical testing of new therapies, those glioblastoma animal models are urgently needed.

Genetically engineered mouse models (GEMMs) have been, so far, the method of choice for identifying the molecular events underlying glioblastoma pathology. Their use has deepened the molecular understanding of glioblastoma initiation and progression and been demonstrated as valuable tools to test new therapies^6^. However, these conventional transgenic approaches are rather time- and cost-intensive. This is mainly due to the extensive intercrossing that is necessary to maintain the required panels of mouse strains.

These difficulties can be avoided by using xenograft or syngraft transplantation models. Indeed, many established human glioblastoma cell lines have been successfully xenotransplanted in immunocompromised mice. However, these cell lines have been propagated *in vitro* for many years in conditions that induce phenotypic changes, such as the use of serum during cultivation^7^. In some cases, it is therefore questionable whether the tumors arising from cell lines are accurately representing the human disease. In recent years the use of patient-derived xenograft (PDX) models has increased drastically. These models have the advantage of developing gliomas that recapitulate the genetic and histological features of the parental tumors and are extremely useful in evaluating specific drug candidates^8^. However, the requirement to use immuno-compromised mice is a major constraint of both PDX and established human cell line xenograft models.

The central importance of the tumor microenvironment in many cancer entities, including glioblastoma, has become indisputably clear in recent years. As it has been shown that human glioblastoma can consist of 30-50% myeloid cells, mainly tumor associated macrophages (TAMs), the role of the immune system in the development and progression of glioblastoma is attracting increased attention^9^. In order to conduct such studies, including the complex interactions of tumor cells with immunosuppressive TAMs and other immune cells such as T cells, it is imperative to establish immunocompetent preclinical models. Notably, the development of new treatment options aiming at affecting/boosting the anti-tumor immune response or to exploit immune cells as therapeutic vehicles, such as chimeric antigen receptor (CAR)-T cells, is completely dependent on models having a functional immune system.

Syngeneic models use the transplantation of murine glioma cells into animals with identical genetic background, and constitute a promising approach to investigate the pathogenic role of tumor-microenvironment interactions. There are a number of established murine glioblastoma cell lines available currently, including GL261, GL26, CT-2A, SMA-560 and 4C8, with GL261 being by far the most extensively utilized^10^. GL261-derived gliomas, however, exhibit a bulky growth pattern^11^, forming a nodular mass without infiltration of tumor cells into the brain parenchyma. Molecularly, GL261 cells are characterized by an activating Kras mutation, which is almost never observed in human GBM^4^. Moreover, these tumors rarely, if ever, exhibit necrosis, which in addition to their invasive growth is a notable characteristic of human glioblastoma.

To address these challenges, we generated a panel of glioblastoma cell lines from neural stem cells and tumor cells of mice with neural stem cell-specific *p53* and *Pten* deletion^12^. Upon orthotopic transplantation into syngeneic hosts, these cell lines formed high-grade gliomas that faithfully recapitulated key characteristics of human glioblastoma. These syngraft tumors displayed cell line-dependent genotypic and phenotypic differences that will make them useful preclinical models to identify and test various treatment approaches, including immunotherapies.

## Results

### Generation of murine glioma cell lines by repeated *in vivo* passaging of *Pten/p53* deleted neural stem cells

Recently, we described a novel genetic model in which tamoxifen-induced neural stem cell (NSC)-specific deletion of *Pten* and *p53* results in the development of brain tumors which were classified as high-grade gliomas (glioblastomas) according to histopathological (necrosis and microvascular proliferation) and molecular features. In line with this classification, the tumors were positive for established glioma markers such as Gfap and Olig2 and show an intense staining for the proliferation marker Ki67^12^. However, this genetic model (hereafter referred to as double knock-out, DKO) exhibits prolonged latency (10 to 24 months after tamoxifen injection), and incomplete penetrance of tumor development (65%), prompting us to generate a faster model based on transplantable murine glioblastoma cells that would be more amenable for experimental work.

We started out with cell lines from DKO mice at two different time points of tumor development:

1. from isolated NSCs two weeks after tamoxifen-induced *pten/p53* deletion (transformed NSC 0; tNSC0). At this time point DKO mice did not show any overt tumor lesions (Figure 1A); however, they all presented with an expansion of the rostral migratory stream (RMS), formed by NSCs that migrate from the sub-ventricular zone (SVZ) of the lateral ventricle (LV) to the olfactory bulb ^12^.
2. from isolated tumor cells of a fully established invasive high grade glioma (murine glioblastoma 0; mGB0) which occurred 12 months after the initial NSCs expansion (Figure 1A)^12^.

**Figure 1.**
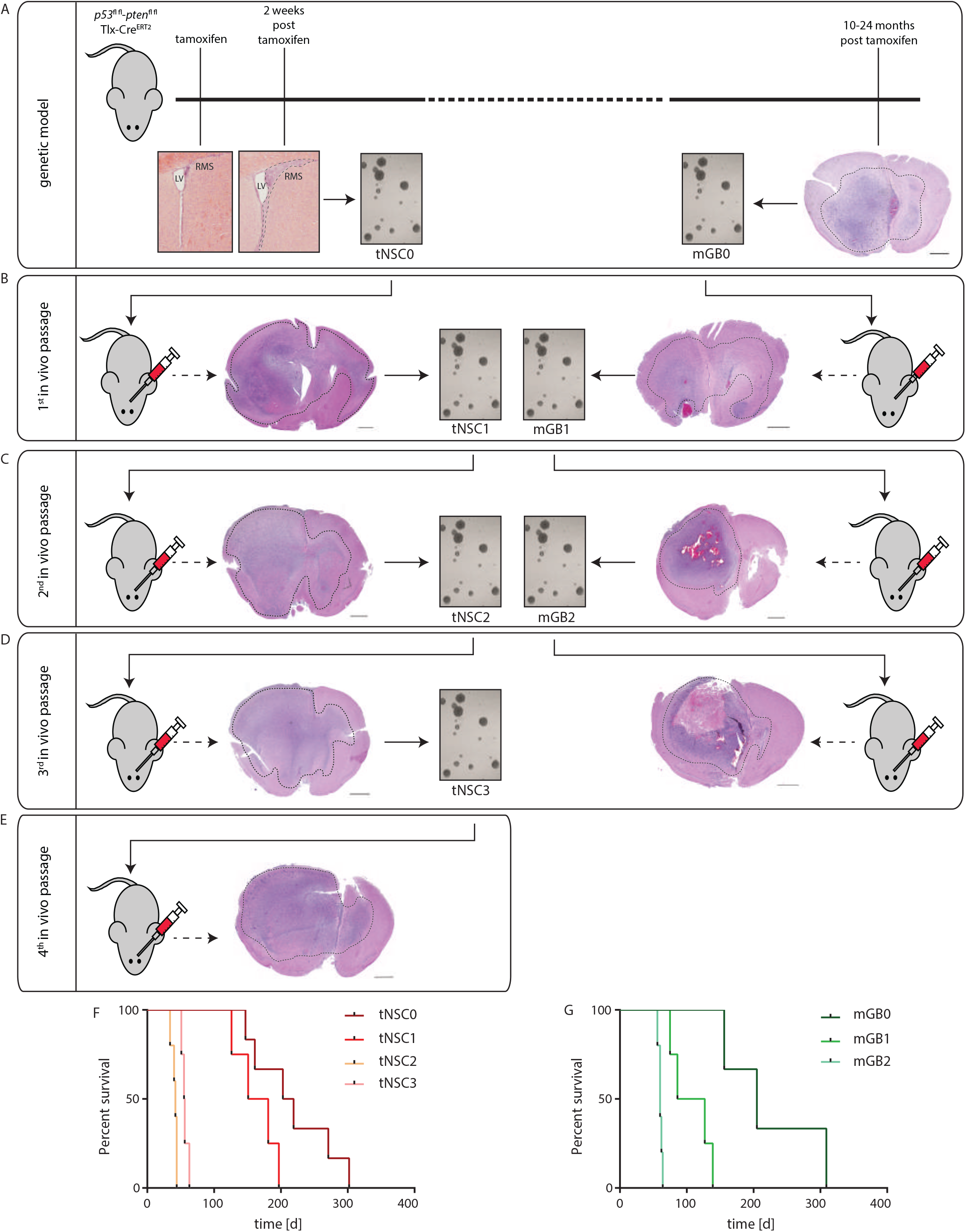
Generation of syngeneic glioma cell lines through *in vivo* passaging of *Pten/p53*-deleted cells isolated from a genetic glioma model. **a** Schematic representation of the *Pten/p53* genetic glioma model from which NSC have been isolated either at early pre-malignant stage (left panel) or from a full blown tumor (right panel). Scale bar = 1000μm Left panel: lateral ventricle (LV) with rostral migratory stream (RMS) at time of tamoxifen injection and at 2 weeks after tamoxifen-induced *Pten/p53* recombination. Sections were stained with hematoxylin and eosin (H&E). Scale bar = 1000μm Right panel: section of a full-blown tumor developed 18 months after tamoxifen-induced *Pten/p53* deletion. Section was stained with hematoxylin and eosin (H&E). Scale bar = 1000μm Dotted line shows tumor area. **b** First *in vivo* passage of tNSC0 (left panel) and mGB0 (right panel). Representative picture of a tNSC0-derived (left panel) and of a mGB0-derived (right panel) glioma. Sections were stained with hematoxylin and eosin (H&E). Scale bars = 1000μm **c** Second *in vivo* passage of tNSC1 (left panel) and mGB1 (right panel). Representative picture of a tNSC1-derived (left panel) and of a mGB1-derived (right panel) glioma. Sections were stained with hematoxylin and eosin (H&E). Scale bars = 1000μm **d** Third *in vivo* passage of tNSC2(left panel) and mGB2 (right panel). Representative picture of a tNSC2-derived (left panel) and of a mGB2-derived (right panel) glioma. Sections were stained with hematoxylin and eosin (H&E). Scale bar = 1000μm **e** Fourth *in vivo* passage of tNSC3 (left panel). Representative picture of a tNSC3-derived (left panel) Sections were stained with hematoxylin and eosin (H&E). Scale bar = 1000μm **f** Kaplan-Meier survival curve of mice transplanted with tNSC0, tNSC1, tNSC2, tNSC3 glioma cell lines. Time = days **g** Kaplan-Meier survival curve of mice transplanted with mGB0, mGB1, mGB2 glioma cell lines. Time = days

In order to promote tumor progression, mGB0 and tNSC0 cells were serially transplanted into immunocompetent C57/Bl6N mice for further two (mGB1, mGB2) and three (tNSC1; tNSC2, tNSC3) *in-vivo* passages, respectively (Figures 1B-E). Upon each re-isolation, the tumor cells were propagated for a few passages in serum-free medium with growth factors and N2 supplement. It has been shown that these culturing conditions avoid serum-induced cell differentiation and keep the tumor cells more similar to the primary tumors from a phenotypic and genomic point of view^7^.

At each *in vivo* passage both tNSC and mGB cell lines gave rise to malignant and highly invasive brain tumors (Figure 1B-E) with 100% penetrance and a progressive shortening of median survival (Figure 1F and G). Remarkably, tNSC1 and mGB1 cells formed glioblastomas with similar latencies and resulted in the death of the animals within 300 days post cell transplantation. This suggests that tNSC1 cells, although isolated from DKO mice at a pre-malignant stage, already have a tumorigenic potential comparable to the mGB0 cells isolated from the fully developed tumors.

All the established glioblastoma cell lines (tNSC0,1, 2 and 3 and mGB0,1 and 2) were propagated in *vitro* for no longer than 10 passages before implantation. To ascertain whether these cells retain their tumorigenicity after longer-term cultivation, tNSC3 cells were kept in serum-free culture, for 60 passages and subsequently injected orthotopically into immunocompetent mice. The survival time of these animals was comparable to that of mice injected with early-passage tNSC3 cells, (Supplemental Figure 1A). In both cases tumor penetrance was 100%. Moreover, tNSC3 cells injected at late passages were still able to give rise to invasive high grade gliomas (Supplemental Figure 1B). These results imply that the tumorigenic potential of our glioblastoma model is not affected by prolonged *in-vitro* culture.

In summary, we showed that NSC-specific *Pten* and *p53* deletion followed by *in-vivo* passaging by orthotopic transplantation resulted in the development of high-penetrance brain tumors within a reasonable time frame.

### Murine glioma cell lines reflect the transcriptome heterogeneity of human glioblastoma

We next used RNA sequencing (RNAseq) to characterize the gene expression profiles of the cultured tNSC (tNSC0-3) and mGB (mGB0-2) cells, along with control non-transformed NSCs (ctrlNSCs).

In order to confirm the genetic background of our tumor cells, we checked for RNAseq reads mapping to the regions of the *Pten* and *p53* genes (Supplementary Figure 2A). As expected, we could not detect any reads corresponding to the *p53* exons 2-10 or *Pten* exon 5 in any of our cell lines (Supplementary Figure 2A) confirming that they are derived from *Pten/p53* double knock-out NSCs. In line with these results, Gene Set Enrichment Analysis (GSEA) analysis of tNSC and mGB cells in comparison to control NSCs suggests a deregulation of p53 pathway-related genes (Supplementary Figure 2B, Supplementary Tables 1,2). Furthermore, we called single nucleotide variants, to confirm that mutations of the IDH1 and IDH2 genes, which are mutated in about 10% of human high-grade gliomas, had not spontaneously occurred during *in-vivo* passaging and tumor development (Supplementary Figure 2C).

Next, we performed multidimensional scaling analysis (MDS) using all genes (Figure 2A). The analysis revealed that tNSC0 and mGB0, which were isolated from the genetic DKO model, still are closely related to the non-transformed NSCs and to each other, even though they were isolated at the beginning and at the end of tumor development. With *in-vivo* passaging, the gene expression profiles diverged, with tNSC- and mGB-derived cell lines appearing to form separate clusters (Figure 2A).

**Figure 2.**
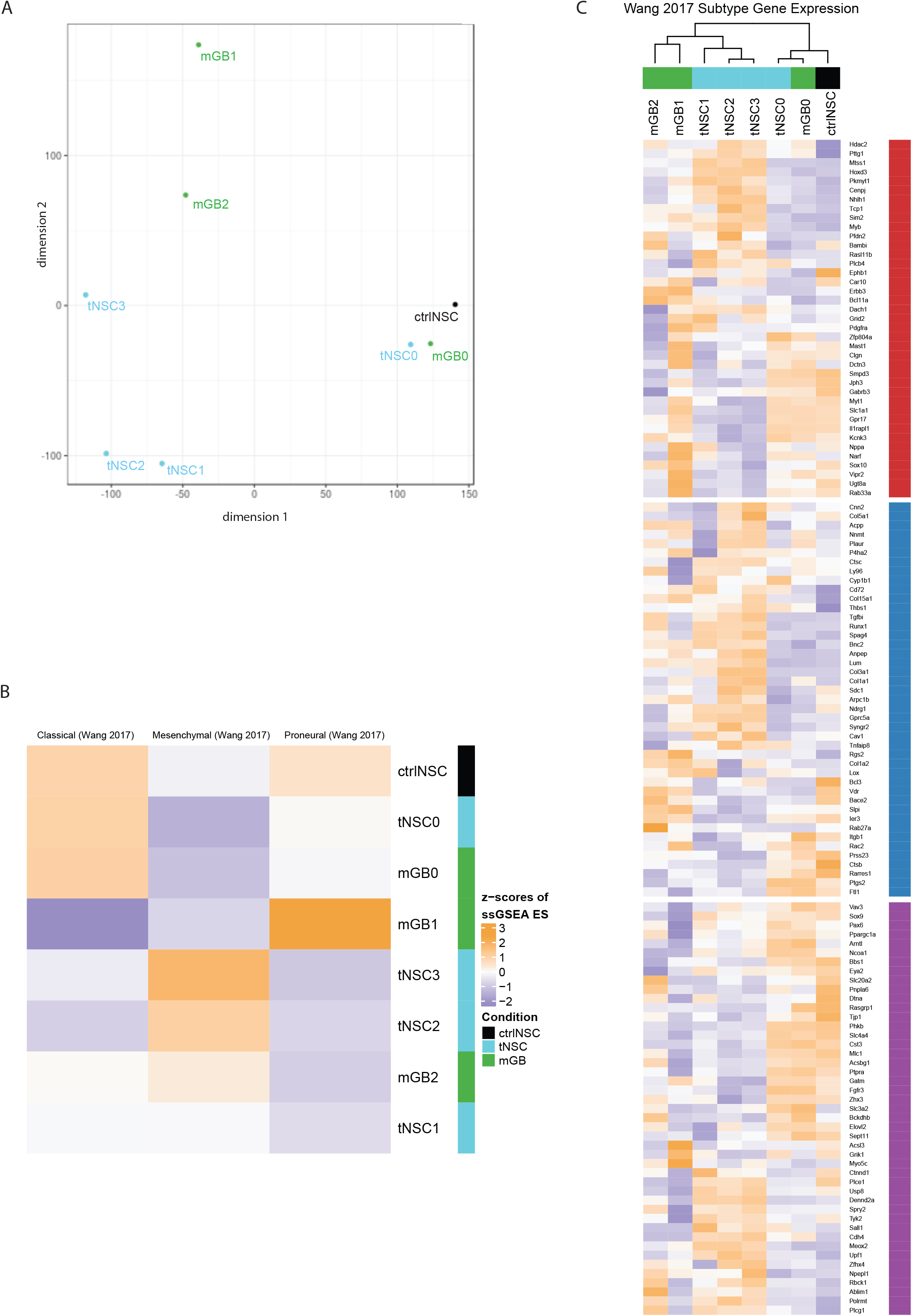
Transcriptomic characterization of the new glioma cell lines finds distinct expression profiles that correspond to known human glioblastoma subtypes. **a** Multidimensional scaling (MDS) analysis of whole transcriptomes from the newly generated glioma cell lines (tNSC0-3, mGB0-2) and of non-transformed NSCs (ctrlNSCs). **b** Heatmap showing z-scores of ssGSEA enrichment scores of the murine glioma cell lines for published glioblastoma subtype gene expression signatures. **c** Heatmap of z-scores of expression of glioblastoma subtype genes, as defined in Wang et al. Genes are labeled with their subtype. The gene expression of cell lines with sample replicates (tNSC4 and ctrlNSC: both n=3) were averaged.

It is commonly accepted that, based on gene expression signatures, IDH^wt^ human glioblastomas can be sub-classified into 3 distinct molecular subtypes: classical, proneural, and mesenchymal^5^. We therefore performed single-sample (ss)GSEA with the expression signatures described in Wang et al.^5^, to relate our cell lines to these well-studied human subtypes (Figure 2B). As expected, control NSCs are enriched for the classical signature, due to the large number of stemness related genes in this set. Consistent with the transcriptome-wide MDS analysis, the mGB0 and tNSC0 cell lines showed similar enrichment patterns to and clustered with control NSCs. Strikingly, the mGB1 cell line showed an increased enrichment score for the proneural signature, while all the other cell lines (tNSC1-3 and mGB2) were relatively enriched for the mesenchymal signature (Figure 2B). *In vivo*, these cell lines showed a progressively aggressive phenotype and decreased survival time of the recipient mice (Figure 1F and G), which is consistent with human glioblastoma, where tumors of the mesenchymal subtype have the poorest prognosis^5, 13, 14^.

We next examined the relative expression of the individual subtype signature genes in greater detail (Figure 2C). Clustering of the cell lines using these genes recapitulated the MDS analysis on all genes (Figure 2A), with the ctrlNSC and early passage samples (tNSC0, mGB0) grouping together and later passages forming distinct clusters based on their lineage. Altogether these results show that the neural stem-cells derived glioblastoma cell lines serially passaged *in vivo* reflect, at the transcriptomic level, the heterogeneity observed in human glioblastoma.

### Characterization of the newly established syngeneic glioblastoma cell lines at the genomic level

In order to characterize the new glioblastoma cell lines at a genomic level we performed whole exome sequencing (WES) and then analyzed copy number aberrations (CNAs) (Supplementary Tables 3-9).

The tNSC0 and mGB0 cell lines showed a limited degree of gross chromosome instability, with the inferred copy number state still being diploid. However, copy number gains and losses acquired after the first *in vivo* passage were maintained in later passages (tNSC1, 2, 3 and mGB1, 2 respectively) (Figure 3). In particular, the cell lines tNSC1-3 showed gross chromosomal alterations such as the gain of a large portion of chromosome 15 and the loss of the remaining part of it, the loss of most of chromosome 14 and gain of chromosome 6.

**Figure 3.**
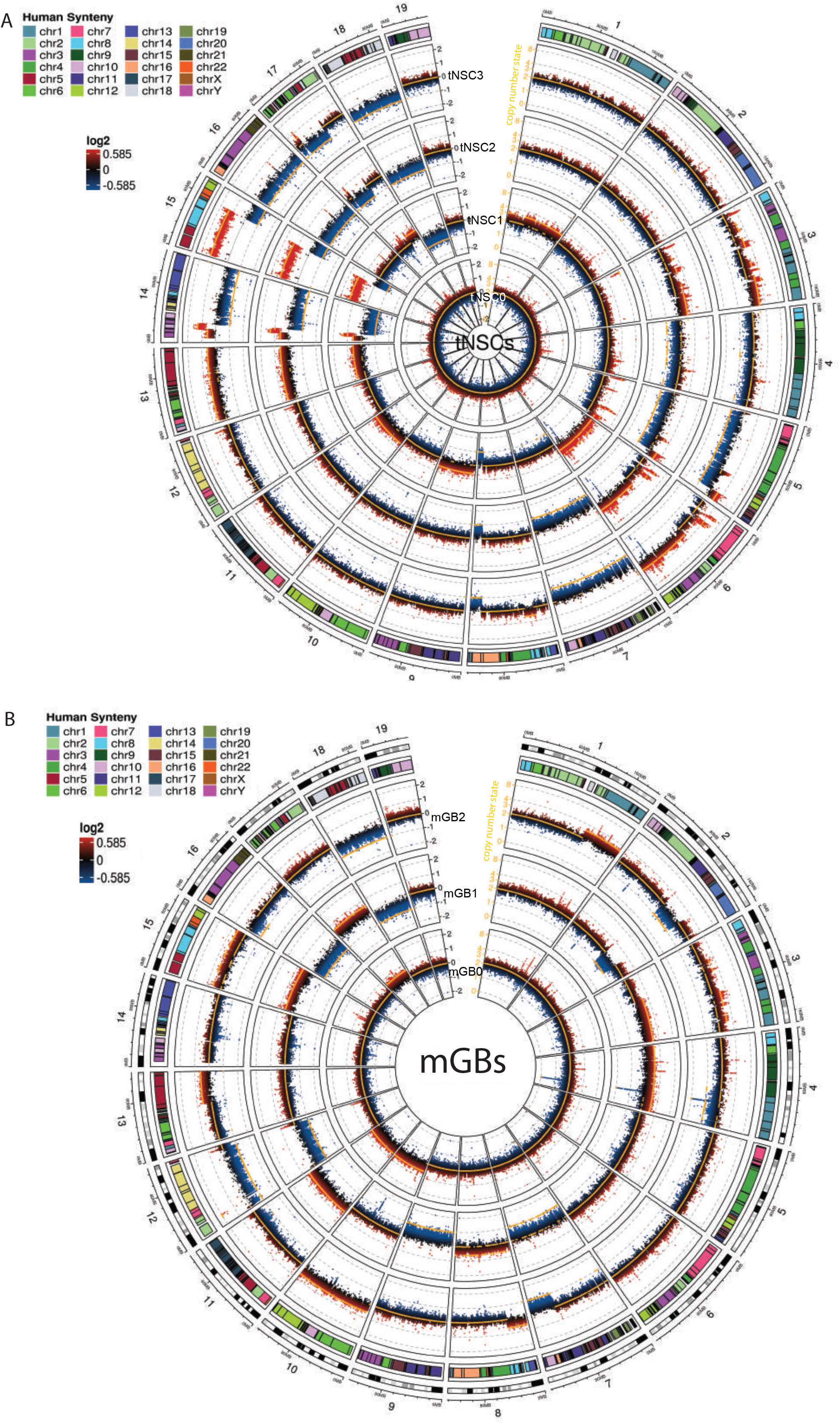
Copy number aberration analysis identifies specific chromosomal aberrations of the new murine glioma cell lines. **a.b** Circos plots showing copy number aberrations (CNA) of tNSCs (a) and mGBs (b) cell lines. CNVkit-calculated log2 ratios for each genomic bin are plotted as coloured points (black y-axis scale; red corresponds to increased log2 ratios, and blue to decreased), with the inferred copy number state plotted as an orange line (orange y-axis scale). Each circular sector represents one chromosome. Human syntenic regions are shown at the top of each chromosome.

Overall, copy number losses were more prevalent than gains. In the tNSC cell lines, for instance, large parts of chromosomes 4, 5, 7, 16 and 18 are lost, and chromosomes 8, 14, 15 and 17 showed focal losses. This result is in line with previous reports on human glioblastoma and other tumor entities where genetic losses are more common than gains ^4, 15^.

In order to relate observed CNAs to known human glioblastoma alterations, we used the Log Fold Change (logFC) values of genes identified via CNA calling to run a pre-ranked GSEA analysis against the human C1 positional gene set MSigDB collection. This analysis allowed us to identify murine chromosomal regions with partial deletions or amplifications that correspond to chromosomal alterations seen in human glioma samples.

In the tNSC1-3 cell lines, a large portion of the gain of the chromosome 15 is syntenic for human chromosome 8. It was recently reported that cell populations that are polyploid for this chromosome are present in human glioblastoma, and that this copy number gain can be used to identify circulating glioblastoma cells^16^. In the same cell lines, a large portion of the mouse chromosome 6 was also gained. This region corresponds to human chromosome 7 and contains an amplification of the Met receptor with a consistently high copy number logFC value (0.56, 1.16 and 1.31 for tNSC1, tNSC2 and tNSC3, respectively). Met is commonly overexpressed or amplified in human glioblastoma. However, in our cell lines, this amplification did not affect the expression levels of Met, suggesting that further genetic and/or epigenetic events might affect Met expression. The tNSC1-3 cell lines also showed a notable deletion in a major part of chromosome 14 including the mouse Rb1 gene that has been reported to be lost or mutated in a subset of human glioblastoma patients^17^. According to the CNAs we detected a partial loss of the mouse Rb1 gene in these cell lines, resulting in a significant (up to 1.6logFC) reduction in Rb1 RNA expression in the tNSC cell lines compared to control NSCs. Furthermore, GSEA performed on the expression data shows a highly significant enrichment of Rb1 regulated genes that were overexpressed in tNSC3 cells in comparison to the control NSCs (Supplementary Figure 2D, Supplementary Table 10). As this gene set represents genes typically downregulated by Rb1^18^, these data suggest that the chr14 deletion contributes to the loss of the tumor suppressive function of Rb1.

The mGB cell lines showed a smaller number of gross genomic alterations, in comparison to the tNSCs. Performing the same pre-ranked GSEA analysis as previously described, we noted a single-copy gain of mouse chromosome 10 in the mGB0 and mGB2 cell lines. This region of the mouse chromosome corresponds to the human 12q13 region, which has been previously reported as amplified in glioblastoma^19^. The Erbb3, Gli1 and Mdm2 genes are all present in this region, and their amplification is known to have effects in human glioblastoma^20, 21, 22^. The amplifications of Erbb3 and Mdm2 correlated with a 6.2 and 1.9 logFC increase in RNA expression of all of these genes, respectively, in mGB cells, compared to control NSCs.

These data illustrate that our murine cell lines harbor genetic alterations known to be important in human glioblastomas. These alterations can lead to changes in RNA expression which may be of functional consequence and thus contribute to the phenotypic differences observed in our cell lines.

### The syngeneic gliomas derived from the newly established cell lines resemble human high-grade gliomas

Upon orthotopic transplantation, all the tNSC and mGB cells lead to the development of high grade gliomas. Although tumor cells were always implanted in the right hemisphere the resulting tumors extended into the contralateral hemisphere (Figure 1B-E), underlining the invasive potential of these tumor cells. In general, tumors obtained from mGBs cells showed intratumoral hemorrhages (Figure 1C, right panel) and extensive necrotic areas (Figure 1D, right panel) in 40% and 15% of the cases, respectively. In comparison 20% and 0% of tNSCs-derived tumors show similar features, indicating that mGBs cells give rise to more malignant high grade gliomas.

The glioblastoma cell lines derived from the last *in-vivo* passages (tNSC3 and mGB2) were characterized more in detail. The cells were stably transduced with GFP to enable visualization and unambiguous identification of tumor cells in tissue samples. Immunohistochemical staining of GFP-expressing tNSC3 and mGB2 cells in formalin fixed paraffin embedded (FFPE) brain sections showed that the tumor margins of the tNSC3- and mGB2-derived gliomas (Figure 4 A and B) were not well delineated due to tumor-cell invasion of the surrounding brain parenchyma (Figure 4C and D), as is characteristic of human glioblastoma.

**Figure 4.**
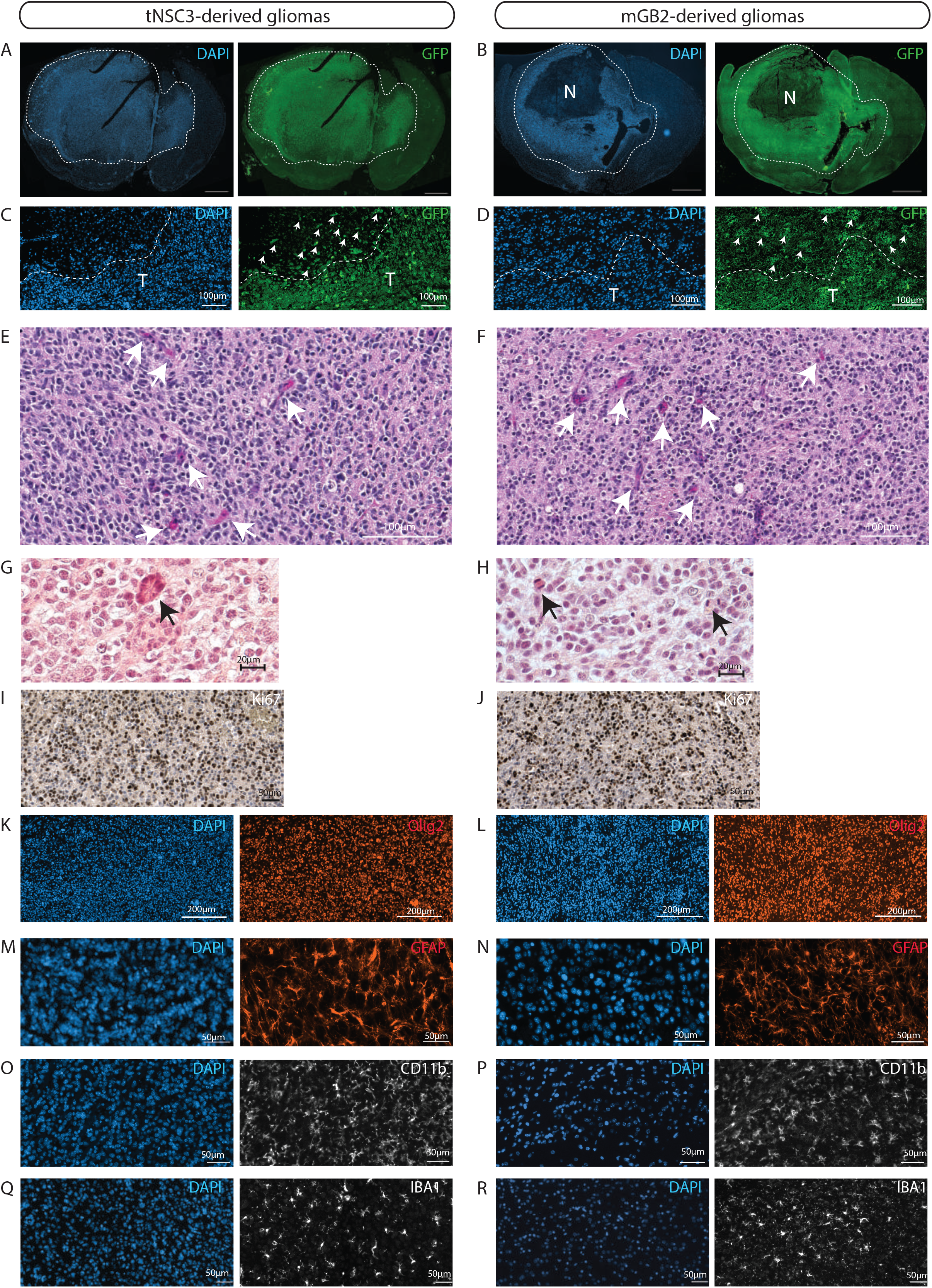
The novel cell lines give rise to orthotopic tumors with characteristics of human glioblastomas. **a,b** GFP-Immunofluorescence staining (green) of tNSC3(a) and mGB2 (b) orthotopic tumors. Cellular nuclei are stained with DAPI and pseudocolored in blue. Scale bars, 1000 μm. Dotted lines show tumor area. N indicates a necrotic area in the mGB2 orthotopic glioma. **c,d** Zoom-in of images from a and b respectively showing area at the tumor border. T indicates tumor area delineated by dotted lines. Arrows denote GFP-positive glioma cells invading the surrounding normal brain parenchyma. **e-h** Histopathological features of orthotopic tNSC3 (e,g) and mGB2 (f,h) gliomas. Sections were stained with hematoxylin and eosin (H&E). Arrows in e,f denote areas of microvascular proliferation; arrows in g,h indicate mitotic figures. **i,j** Ki67 expression (purple) in orthotopic tNSC3 (e) and mGB2 (f) gliomas detected by immunohistochemistry. The sections were counterstained with hematoxylin. **k-r** Immunofluorescence staining for Olig2 (k,l), Gfap (m,n), Cd11b (o,p), Iba1 (q,r) within tumor areas from orthotopic tNSC3 (I,k,m,o) and mGB2 (j,l,n,p) gliomas. Cellular nuclei are stained with DAPI and pseudocolored in blue.

In addition, these tumors had histopathological features of high-grade gliomas, such as microvascular proliferation and the presence of mitotic figures (Figure 4 E-H, Supplemental Figure 1C,D). Furthermore, tNSC3- and mGB2-derived tumors were strongly positive for the proliferation marker Ki67 (Figure 4I and J) and for the human glioma markers Olig2 (Figure 4K and L) and Gfap (Figure 4 M and N). Previously, expression of branched-chain amino acid transaminase 1 (BCAT1) was found to be exclusive to human gliomas with wild-type IDH1 and IDH2^23^. In agreement with this, immunohistochemical analysis of our tumors revealed prominent staining for Bcat1 (Supplementary Figure 2E).

Next, we examined the tumor microenvironment with a focus on immune cells. Staining of tNSC3 and mGB2 tumors for Cd11b and Iba1 indicated a strong presence of myeloid cells (Figure 4O-R) while staining for Cd3 identified very few lymphocytes (data not shown). These results are in accordance with human high-grade gliomas where it has been shown that tumor–associated macrophages (TAMs) can constitute up to 30% of the tumor mass^9^ but are also immunologically “cold tumors” due to the very low number of tumor-infiltrating lymphocytes (TILs)^24^.

Altogether, these data show that the tumors formed after transplantation of the new NSC-derived mouse cell lines resemble human glioblastoma with respect to histopathological and molecular features and to immune cell infiltration.

## Discussion

Despite recent advances in their characterization at genomic, epigenomic and transcriptomic levels^4, 5, 25, 26^, glioblastomas remain almost invariably deadly tumors with very limited treatment options. Therapeutic progress is decidedly impeded by the tumors’ genotypic and phenotypic heterogeneity and complexity of tumor-microenvironment interactions. Likely reasons for the failure of vaccination, chimeric antigen receptor, adoptive T-cell and immune checkpoint inhibitor approaches are glioblastomas’ low mutational burden, subclonal heterogeneity and the highly suppressive tumor microenvironment, facilitated by the tumor-associated macrophages that hamper T cell responses. Attempts to inhibit or reprogram cells of the tumor microenvironment, such as TAMs, have invariably failed in clinical trials, despite promising results in murine models^1^. These failures to predict the success of clinical trials using current preclinical animal models evidence the strong need to develop immunocompetent glioblastoma models that faithfully recapitulate the human disease.

The subventricular zone (SVZ) of the adult human brain contains a population of neural stem cells (NSCs), which are capable of multilineage differentiation^27, 28^. Mutations of key tumor suppressors in SVZ cells have been shown to initiate the development of glioblastomas of diverse phenotypes^29, 30^. We previously described a genetic mouse model with NSC-specific knockout of *Pten* and *p53* genes^12^, driven by the Tlx promoter^31^. This genetic mouse model is based on the inactivation of two tumor suppressors relevant to glioblastoma specifically in the stem cell population that is thought to commonly give rise to these tumors. Moreover, it retains the differentiation potential to produce the range of genotypic and phenotypic heterogeneity that is observed in human glioblastoma. Following up to three *in vivo* passages of the SVZ NSCs and tumor cells, we were able to generate cell lines that, after syngeneic transplantation, reproducibly formed high-grade gliomas displaying key characteristics of human glioblastoma. Transcriptome analysis of the cells revealed cell line-specific enrichment of RNA expression signatures^5^ of the proneural, classical and mesenchymal human glioblastoma subtypes. This suggests that, despite their common cell of origin, the cell lines through *in-vivo* passaging have diversified to present a level of heterogeneity resembling that of human glioblastoma. In line with these findings, genome analysis identified copy number aberrations that were conserved through *in-vivo* passaging, including some that correspond to known chromosome aberrations of human glioblastoma. These data show that the generated mouse glioblastoma cell lines mirror human glioblastoma genetic heterogeneity.

One of the most striking and clinically relevant characteristics of human glioblastoma is their highly invasive growth. Glioblastoma cells typically infiltrate large regions of the brain including the brain parenchyma, the corpus callosum and the contralateral hemisphere. This invasive growth constitutes one of the biggest clinical challenges, preventing complete surgical resection and almost invariably resulting in tumor recurrence. The mouse glioblastomas resulting from the orthotopic transplantation of our syngeneic lines display remarkably similar infiltrative growth patterns as the tumors invaded the contralateral hemisphere in all cases. Furthermore, they showed necrotic areas and microvascular proliferation, both defining characteristics of human glioblastoma. In contrast, the currently prevalently utilized syngeneic mouse glioblastoma model, the cell line GL261, is lacking most of these features and presents with bulky tumor growth and no obvious infiltration of the adjacent normal brain tissue^11^.

Human glioblastoma are considered immunologically “cold” tumors that almost completely lack observable infiltration by lymphocytes^24^. Nevertheless, the tumor bulk often contains large numbers of tumor-associated macrophages with typical immunosuppressive phenotype. These myeloid cells have been proposed to be responsible for the severe suppression of the tumor infiltrating lymphocyte population and thus to render most novel immunotherapeutic strategies ineffective in the case of glioblastoma^1^. In order to further optimize such therapies and design immune-altering drugs that can elicit an anti-tumor response, the use of immunocompetent glioblastoma models, in addition to patient-derived *in vitro* glioblastoma organoid cultures ^32^ and the commonly used patient-derived xenograft models, is an absolute necessity. Our mouse glioblastoma model shows extensive myeloid-cell infiltration with barely any tumor infiltrating lymphocytes in the tumor mass. These observations strongly suggest that the immunosuppressive environment of the human glioblastoma is recapitulated by our transplantation model.

One of the main strengths of our panel of syngeneic glioblastoma cell lines is their full transcriptomic and genomic characterization. This will enable researchers to extract information about their genes of interest and choose the appropriate cellular model for their studies. Moreover, additional genetic modifications of the cell lines *in vitro*, i.e. by CRISPR/Cas9 or viral transduction, are easily possible. The usefulness of our transplantation models was already demonstrated in a current study about the role of the SOX10 master regulator transcription factor in the determination of glioblastoma transcriptional and phenotypic transitions (Wu et al., submitted; manuscript enclosed). In this study, the mGB1 syngeneic, orthotopic transplantation model was used to show that, upon suppression of Sox10, syngraft tumors not only acquired a more aggressive growth phenotype, resulting in significantly worse survival of the animals, but also showed a much greater extent of infiltration by myeloid cells. These results demonstrate that our panel of models can successfully be utilized in deciphering determinants of glioblastoma malignancy and, thus, contributes to future efforts in targeting this disease.

In summary, our model allows researchers to simply and reproducibly generate syngeneic mouse glioblastomas that can easily be further genetically modified. The generated tumors represent faithful models of human glioblastoma on histopathological and molecular levels in a fully immunocompetent background. Due to the varying malignancies of our cell lines in the animals, manipulations of tumor aggressiveness, testing of immunomodulatory therapies, as well as long-term treatment options with wide therapeutic windows will be more easily possible than with many current animal models. These characteristics make our syngeneic transplantation model an attractive, unique and valuable tool for basic and translational glioblastoma research.

## Methods

### Animals

*Tlx*-CreER^T2^/*p53*-floxed/*pten*-floxed (double knockout [DKO]) mice were described in^12^. To induce recombination of floxed alleles, 4-week-old mice were injected intraperitoneally with 1 mg tamoxifen (S5007 Sigma) in 5% ethanol and 95% oil (T5648 Sigma) for 5 consecutive days. Animal experiments were approved by the German responsible authority (Regierungspräsidium Karlsruhe) and performed in conformity with the German law for Animal Protection (animal license number: G-156/15, G-199/11).

### Cell isolation

NSCs were isolated from the subventricular zone (SVZ) of mice 2 weeks after the injection of either tamoxifen or oil according to^33^. For tumor cell isolation, glioma-bearing mice were euthanized with carbon dioxide; brains were minced and dissociated in Leibovitz-L15 (Life Technologies) containing 10 U/mL papain, 5 mM EDTA, and 200 U/mL DNAse. Cells were grown according to^33^.

### Cell transduction

Lentiviral transduction with a construct encoding eGFP (Plasmid #14883, Addgene) was performed in order to label the cells. For virus production one 10 cm dish HEK293T cells was transfected with 8 μg target vector; 4 μg psPAX2; 2 μg pVSVg and 42 μg polyethylenimine (Alfa Aesar). HEK293T cells were cultivated in N2-supplemented serum-free medium. Virus-containing medium was transferred from HEK293T cells to the target cells and replaced by cultivation medium after 24 h.

### Orthotopic intracranial injections

Mice were anesthetized with isoflurane and placed on a stereotaxic frame. A total of 5 × 10^5^ cells in 2 μL phosphate-buffered saline were injected 2 mm lateral (right) to the bregma and 3 mm deep at a flow of 0.2 μL/min using a 10-μL precision microsyringe (World Precision Instruments, Inc) with a 34G needle.

### Image acquisition

Pictures were captured with Zeiss Axio-Scan.Z1 using ZEN software (Zeiss) or with the MEA53100 Eclipse Ti-E inverted microscope (Nikon) using MQS31000 NIS-ELEMENTS AR software (Nikon) with camera MQA11550 DS-Qi1MC for bright field images and MQA11010 DS-Fi1 for immunofluorescent images.

### Immunohistochemistry and immunofluorescence staining

Formalin fixed paraffin-embedded sections (5 μm) were stained as described previously^34^ using the following primary antibodies: rabbit anti-Ki67 (ab15580, Abcam), rabbit anti-OLIG2 (ab109186, Abcam), rabbit anti CD11b (ab133357, Abcam), rabbit anti Iba1 (019-1974, Wako), chicken anti GFP (ab1397, Abcam), mouse anti Bcat1 (TA504360, OriGene), mouse anti-GFAP (644701, BioLegend). Secondary antibodies were: goat anti-rabbit Alexa Fluor 546 (A11071 Thermo Fischer Scientific) and goat anti-mouse Alexa Fluor 555 (A21422, Thermo Fischer Scientific). For immunohistochemistry, the biotinylated goat anti-rabbit immunoglobulin-G (BA1000) were used together with DAB peroxidase (horseradish peroxidase) substrate kit (SK-4100), both from Vector Laboratories.

### Library preparation and target enrichment of genomic DNA for whole exome DNA sequencing

Genomic DNA was fragmented to a size range of 150-250 bp using the Covaris S220 instrument. The fragmented DNA was end repaired and adenylated. The SureSelect Adaptor was then ligated followed by a pre-amplification. After library preparation target regions were captured by hybridization of biotinylated baits (library probes) using Agilent SureSelect XT Human All Exon V6. Captured target sequences were then isolated using streptavidin coated magnetic beads. Subsequently the appropriate 8 bp single index tags were added during sequencing library amplification. All steps were done using Agilent SureSelect XT Reagent Kit. Final quality control was done using Qubit 3 fluorometer and Agilent Bioanalyzer 2100. The libraries were sequenced in paired-end mode (2 × 50 nt) on a NovaSeq6000 S2 Flowcell resulting in ~200 million distinct sequencing reads per library.

### mRNA-focused library preparation of total RNA for NGS

Poly(A)-positive mRNA transcripts were isolated from the total RNA by binding to magnetic oligo(d)T beads (RNA purification beads). After elution from the beads and fragmentation, the mRNA was reverse transcribed during first and second strand synthesis yielding double-stranded cDNA. The ends of the cDNA molecules were end repaired and an adenosine overhang was added to all 3’ ends (A-tailing step) to facilitate the ligation of the adapter molecules. These adapters have individual dedicated index sequences and add the Illumina specific sequences that are needed for amplification, flow cell hybridization and sequencing. 8 bp single-index NEXTflex DNA barcodes (Bioo Scientific) were used. The final ligation product was amplified via PCR. Double-stranded cDNA sequencing libraries were further checked for quality and quantity using Qubit 3 fluorometer and Agilent Bioanalyzer 2100. The libraries were sequenced in paired-end mode (2 × 50 nt) on a NovaSeq6000 S2 Flowcell resulting in ~50 million distinct sequencing reads per library.

### Bioinformatic data analysis

#### WES data – basic processing

Whole exome sequencing paired end reads in fastq format were aligned using *BWA* (v0.7.15) to the mm10 reference genome with options: -M ‒T 0. Duplicates were marked using *sambamba* (0.6.6).

#### WES – CNV analysis

CNV analysis was performed using *CNVkit* (v0.9.6)^35^. Tumor WES samples (tNSC1-4, GBM1-3) were analyzed with reference to the two normal WES samples, ctrlNSC and normal splenocytes. The mm10 genome .fasta was used to calculate accessibility as follows: access mm10.fa -s 10000 ‒o mm10_accessibility.bed. The S0276129 bedfile with Agilent capture array loci, lifted over to the mm10 genome, was used as the target regions. Regions were annotated using UCSC’s mm10 flat reference (http://hgdownload.soe.ucsc.edu/goldenPath/mm10/database/refFlat.txt.gz, accessed 21 November 2019).

Calling was performed with the batch command and the following options: batch TUMOR_SAMPLES --normal NORMAL_SAMPLES --targets S0276129_mm10.bed --fasta mm10.fa --access mm10_accessibility.bed --annotate refFlat.txt --drop-low-coverage. Copy number state segmentation was performed using the ‘cbs’ method as implemented in the R package *DNAcopy* (v1.54.0).

The results of the CNV calling were visualized using the R package *circlize* (v0.4.4)^36^in *R* v3.5.1. Regions overlapping annotated segmental duplications (downloaded from the UCSC Table Browser, accessed 25 January 2020) were removed before plotting due to potential issues with inferring copy number states in these repetitive regions. For ease of visualization, extreme values with *CNVkit* log2 copy ratio < −2 or > 2 were set to this minimum and maximum. The integer *CNVkit* segmented copy number states were smoothed using the rollarray command from *zoo* (v1.8-2) with the options width = 301, partial=TRUE.

The identification of human syntenic regions with observable CNAs in the tNSC and mGB cell lines was done using GSEA against the human “c1.all.v7.0.symbol” dataset with gene names adapted to mouse. The analysis was performed as a pre-ranked test using logFC CNA values produced in the CNA calling.

#### RNA-seq data – basic processing

RNA-seq reads in .fastq format were aligned with *STAR* (v2.5.2a)^37^ to the mm10 reference genome and the Gencode M2 reference transcriptome, using the following options: -- outFilterMismatchNmax 2 --outFilterMismatchNoverLmax 0.05 --alignIntronMax 1 -- outFilterMatchNminOverLread 0.95 --outFilterScoreMinOverLread 0.95 --outFilterMultimapNmax 10 --outFilterIntronMotifs RemoveNoncanonical --outFilterType BySJout --outSAMunmapped Within --outSAMattributes Standard --alignIntronMin 21 --outFilterMatchNmin 16. Overlapping read pairs were clipped using the *clipOverlap* tool in *bamUtil* (v1.0.9) ^38^. The raw counts matrix was generated using *featureCounts* (v1.5.3)^39^ using options -Q 255 -p -t exon against the M2 transcriptome. Aligned data were quality controlled using *RSeQC* (v2.6.4)^40^.

#### RNA-seq - transcriptome analysis

This analysis was performed in *R* (v3.4.3). The raw counts matrix produced by *featureCounts* was pre-filtered to remove genes with low expression, keeping those with >10 reads in >2 samples, and normalized using the *limma-voom* (v3.34.4)^41^ function *voomWithQualityWeights*. Gencode M2 Ensembl Gene IDs were mapped to other identifiers using *org.Mm.eg.db* (v3.5.0). Mouse gene identifiers were mapped to their human homologues using the Jackson Laboratory HomoloGene table (http://www.informatics.jax.org/downloads/reports/HOM_MouseHumanSequence.rpt, accessed 25 March 2019).

Gene Set Enrichment Analysis was performed for each condition against the Wang 2017 subtype classifier^42^. Conditions with multiple replicates (ctrlNSC and tNSC4) were averaged, and then each was tested against the 50 subtype genes from the Wang classification, mapped to mouse homologues as described above, using the *ssgsea* option in *GSVA* (v1.26.0)^43^. MDS plots were generated using the complete expression matrix, with replicates averaged as above.

Heatmaps were visualized using *ComplexHeatmap* (v1.17.1)^44^ and other graphics using *ggplot2* (v2.2.1). The rows/columns in both heatmaps in Figure 3 were clustered with the Ward.D2 method using 1-Pearson’s correlation as the distance metric.

#### WES/RNA-seq – Idh1/2 hotspot mutation analysis

Single nucleotide variants (SNVs) were called in the WES and RNA-seq data using *BCFTools* (v1.9)^45^ at the 3 hotspots (Idh1: R132 = 1:65170977-65170979; Idh2: R140 = 7:80099112-80099114; R172 = 7:80099016-80099018). *bcftools mpileup* was used on the full set of samples for each data type, and consensus calling performed with *bcftools call ‒c*.

The GSEA for the enrichment in the p53 and rb1 transcriptional signatures was performed as a pre-ranked analysis of the genes differentially expressed between tNSC and mGB cell lines and the ctrlNSCs. Only genes with FDR<0.01 were considered for the analysis. The expression signatures used were MSigDb “h.all.v7.0.symbols” and “c2.all.v7.0.symbols” adapted for mouse gene names, respectively.

## Data availability

RNA seq and WES data have been deposited at the GEO; accession number GSE145559.

## Acknowledgments

The authors thank Christine Bauer and Angelika Krischke for contributing their technical expertise to this manuscript. The authors also thank the DKFZ Central Animal Laboratory for animal care, the DKFZ Light Microscopy Facility for assistance in imaging, and the DKFZ Flow Cytometry Facility for cell sorting. This work was supported by the Helmholtz Alliance Preclinical Comprehensive Cancer Center (P.A.).

## Ethics declarations

**Supplementary Figure 1.**
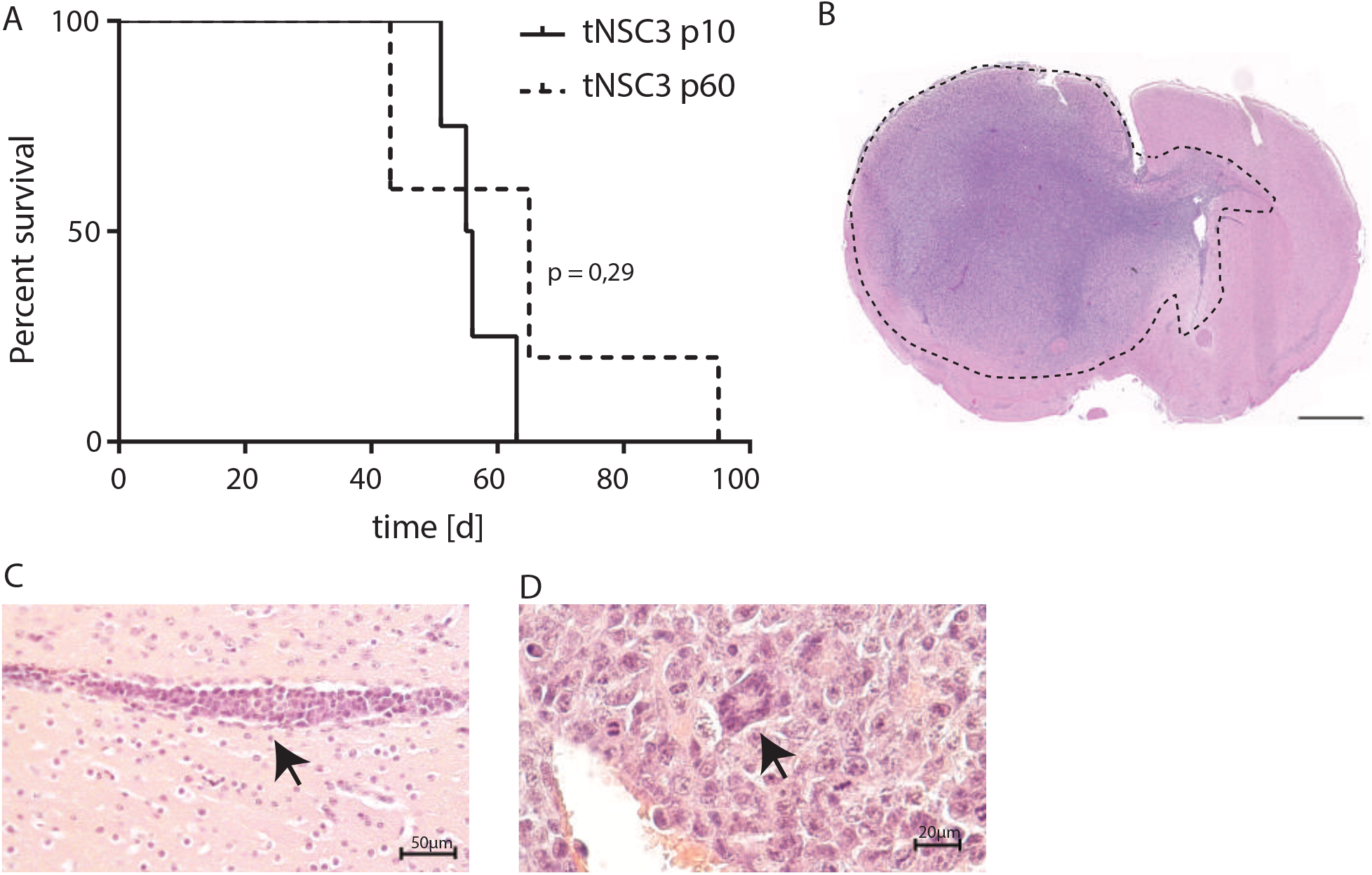
**a** Kaplan-Meier survival curve of mice transplanted with tNSC3 cells at early (p10) and late passages (p60). Time = days. Statistical analysis: Mantel-Cox test **b** Representative picture of a tNSC3 orthotopic glioma derived from cells at late passage (p60). Section was stained with hematoxylin and eosin (H&E). Scale bar = 1000μm. Tumor area is delineated by a dotted line. **c,d** Histopathological features of orthotopic tNSC3 late passage gliomas. Sections were stained with hematoxylin and eosin (H&E). Arrow in c denotes areas of perivascular growth; arrow in d indicates a mitotic figure.

**Supplementary Figure 2.**
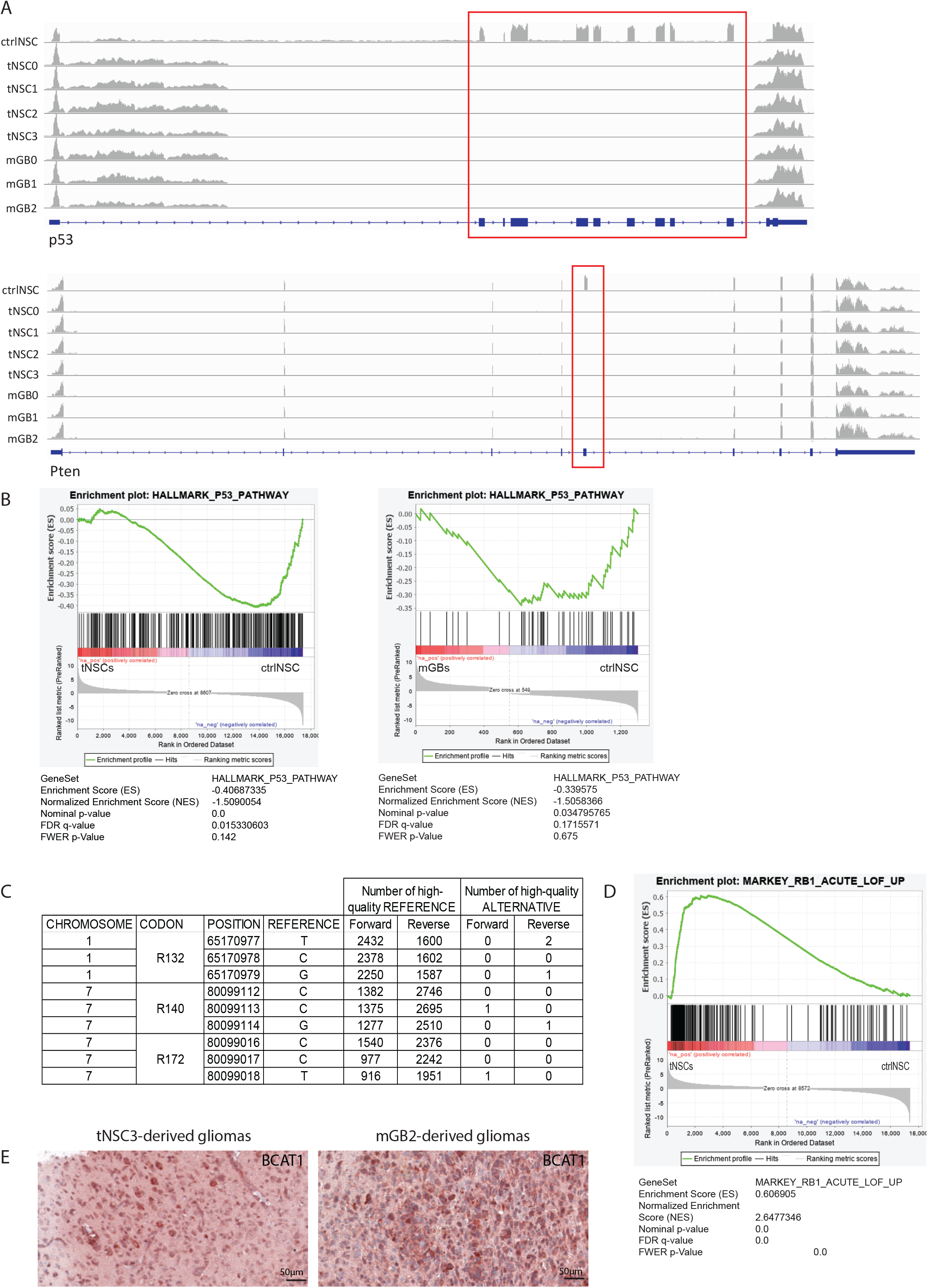
**a** Schematic representation of RNA sequencing reads mapping on p53 and Pten. Blue boxes represent exons. The red rectangles highlight the lack of coverage in exons 2-10 for p53 and exon 5 for Pten in the tNSC and GBM cell lines. These exons are deleted in the DKO genetic mouse model following tamoxifen-induced cre-mediated recombination. **b** Gene Set Enrichment Analysis plot for the mSigDB Hallmark pathway “p53-regulated genes” in tNSCs (left panel) and GBMs (right panel) cell lines, compared to ctrlNSCs. **c** Table showing RNA transcript reads and genotype for *Idh1* and *Idh2* mutation-specific codons that correspond to the hotspots commonly mutated in human glioma samples. **d** Gene Set Enrichment Analysis plot for mSigDB “Markey_RB1_Acute_LOF_UP” in tNSCs cell lines versus ctrlNSCs. **e** Bcat1 expression (red) in orthotopic tNSC3 (left panel) and mGB2 (right panel) gliomas detected by immunohistochemistry. The sections were counterstained with hematoxylin.

## References

1. Sampson JH, Gunn MD, Fecci PE, Ashley DM. Brain immunology and immunotherapy in brain tumours. Nat Rev Cancer 20, 12–25 (2020).

2. Louis DN, et al. The 2016 World Health Organization Classification of Tumors of the Central Nervous System: a summary. Acta Neuropathol 131, 803–820 (2016).

3. Weller M, et al. European Association for Neuro-Oncology (EANO) guideline on the diagnosis and treatment of adult astrocytic and oligodendroglial gliomas. Lancet Oncol 18, e315–e329 (2017).

4. Brennan CW, et al. The somatic genomic landscape of glioblastoma. Cell 155, 462–477 (2013).

5. Wang Q, et al. Tumor Evolution of Glioma-Intrinsic Gene Expression Subtypes Associates with Immunological Changes in the Microenvironment. Cancer Cell 32, 42–56.e46 (2017).

6. Miyai M, Tomita H, Soeda A, Yano H, Iwama T, Hara A. Current trends in mouse models of glioblastoma. J Neurooncol 135, 423–432 (2017).

7. Lee J, et al. Tumor stem cells derived from glioblastomas cultured in bFGF and EGF more closely mirror the phenotype and genotype of primary tumors than do serum-cultured cell lines. Cancer Cell 9, 391–403 (2006).

8. Patrizii M, Bartucci M, Pine SR, Sabaawy HE. Utility of Glioblastoma Patient-Derived Orthotopic Xenografts in Drug Discovery and Personalized Therapy. Front Oncol 8, 23 (2018).

9. Graeber MB, Scheithauer BW, Kreutzberg GW. Microglia in brain tumors. Glia 40, 252–259 (2002).

10. Oh T, et al. Immunocompetent murine models for the study of glioblastoma immunotherapy. J Transl Med 12, 107 (2014).

11. Chen X, et al. RAGE expression in tumor-associated macrophages promotes angiogenesis in glioma. Cancer Res 74, 7285–7297 (2014).

12. Costa B, et al. Intratumoral platelet aggregate formation in a murine preclinical glioma model depends on podoplanin expression on tumor cells. Blood Adv 3, 1092–1102 (2019).

13. Klughammer J, et al. The DNA methylation landscape of glioblastoma disease progression shows extensive heterogeneity in time and space. Nat Med 24, 1611–1624 (2018).

14. Wang L, et al. The Phenotypes of Proliferating Glioblastoma Cells Reside on a Single Axis of Variation. Cancer Discov 9, 1708–1719 (2019).

15. Beroukhim R, et al. The landscape of somatic copy-number alteration across human cancers. Nature 463, 899–905 (2010).

16. Gao F, et al. Circulating tumor cell is a common property of brain glioma and promotes the monitoring system. Oncotarget 7, 71330–71340 (2016).

17. Goldhoff P, et al. Clinical stratification of glioblastoma based on alterations in retinoblastoma tumor suppressor protein (RB1) and association with the proneural subtype. J Neuropathol Exp Neurol 71, 83–89 (2012).

18. Eguchi T, Takaki T, Itadani H, Kotani H. RB silencing compromises the DNA damage-induced G2/M checkpoint and causes deregulated expression of the ECT2 oncogene. Oncogene 26, 509–520 (2007).

19. Rao SK, Edwards J, Joshi AD, Siu IM, Riggins GJ. A survey of glioblastoma genomic amplifications and deletions. J Neurooncol 96, 169–179 (2010).

20. Duhem-Tonnelle V, et al. Differential distribution of erbB receptors in human glioblastoma multiforme: expression of erbB3 in CD133-positive putative cancer stem cells. J Neuropathol Exp Neurol 69, 606–622 (2010).

21. Santoni M, et al. Essential role of Gli proteins in glioblastoma multiforme. Curr Protein Pept Sci 14, 133–140 (2013).

22. Biernat W, Kleihues P, Yonekawa Y, Ohgaki H. Amplification and overexpression of MDM2 in primary (de novo) glioblastomas. J Neuropathol Exp Neurol 56, 180–185 (1997).

23. Tonjes M, et al. BCAT1 promotes cell proliferation through amino acid catabolism in gliomas carrying wild-type IDH1. Nat Med 19, 901–908 (2013).

24. Gajewski TF, Schreiber H, Fu YX. Innate and adaptive immune cells in the tumor microenvironment. Nat Immunol 14, 1014–1022 (2013).

25. Ceccarelli M, et al. Molecular Profiling Reveals Biologically Discrete Subsets and Pathways of Progression in Diffuse Glioma. Cell 164, 550–563 (2016).

26. Sturm D, et al. Hotspot mutations in H3F3A and IDH1 define distinct epigenetic and biological subgroups of glioblastoma. Cancer Cell 22, 425–437 (2012).

27. Doetsch F, Caille I, Lim DA, Garcia-Verdugo JM, Alvarez-Buylla A. Subventricular zone astrocytes are neural stem cells in the adult mammalian brain. Cell 97, 703–716 (1999).

28. Sanai N, et al. Unique astrocyte ribbon in adult human brain contains neural stem cells but lacks chain migration. Nature 427, 740–744 (2004).

29. Alcantara Llaguno S, et al. Malignant astrocytomas originate from neural stem/progenitor cells in a somatic tumor suppressor mouse model. Cancer Cell 15, 45–56 (2009).

30. Lee JH, et al. Human glioblastoma arises from subventricular zone cells with low-level driver mutations. Nature 560, 243–247 (2018).

31. Liu HK, et al. The nuclear receptor tailless induces long-term neural stem cell expansion and brain tumor initiation. Genes Dev 24, 683–695 (2010).

32. Jacob F, et al. A Patient-Derived Glioblastoma Organoid Model and Biobank Recapitulates Inter- and Intra-tumoral Heterogeneity. Cell 180, 188–204 e122 (2020).

33. Guo W, Patzlaff NE, Jobe EM, Zhao X. Isolation of multipotent neural stem or progenitor cells from both the dentate gyrus and subventricular zone of a single adult mouse. Nat Protoc 7, 2005–2012 (2012).

34. Peterziel H, et al. Expression of podoplanin in human astrocytic brain tumors is controlled by the PI3K-AKT-AP-1 signaling pathway and promoter methylation. Neuro Oncol 14, 426–439 (2012).

35. Talevich E, Shain AH, Botton T, Bastian BC. CNVkit: Genome-Wide Copy Number Detection and Visualization from Targeted DNA Sequencing. PLoS Comput Biol 12, e1004873 (2016).

36. Gu Z, Gu L, Eils R, Schlesner M, Brors B. circlize Implements and enhances circular visualization in R. Bioinformatics 30, 2811–2812 (2014).

37. Dobin A, et al. STAR: ultrafast universal RNA-seq aligner. Bioinformatics 29, 15–21 (2013).

38. Jun G, Wing MK, Abecasis GR, Kang HM. An efficient and scalable analysis framework for variant extraction and refinement from population-scale DNA sequence data. Genome Res 25, 918–925 (2015).

39. Liao Y, Smyth GK, Shi W. featureCounts: an efficient general purpose program for assigning sequence reads to genomic features. Bioinformatics 30, 923–930 (2014).

40. Wang L, Wang S, Li W. RSeQC: quality control of RNA-seq experiments. Bioinformatics 28, 2184–2185 (2012).

41. Law CW, Chen Y, Shi W, Smyth GK. voom: Precision weights unlock linear model analysis tools for RNA-seq read counts. Genome Biol 15, R29 (2014).

42. Wang Q, et al. Tumor Evolution of Glioma-Intrinsic Gene Expression Subtypes Associates with Immunological Changes in the Microenvironment. Cancer Cell 33, 152 (2018).

43. Hanzelmann S, Castelo R, Guinney J. GSVA: gene set variation analysis for microarray and RNA-seq data. BMC Bioinformatics 14, 7 (2013).

44. Gu Z, Eils R, Schlesner M. Complex heatmaps reveal patterns and correlations in multidimensional genomic data. Bioinformatics 32, 2847–2849 (2016).

45. Li H. A statistical framework for SNP calling, mutation discovery, association mapping and population genetical parameter estimation from sequencing data. Bioinformatics 27, 2987–2993 (2011).

